# Genetic and environmental contributions to eigengene expression

**DOI:** 10.1101/2024.05.13.593914

**Authors:** Nathan A Gillespie, Tyler R. Bell, Gentry C Hearn, Jonathan L. Hess, Ming T. Tsuang, Michael J. Lyons, Carol E. Franz, William S. Kremen, Stephen J. Glatt

**Affiliations:** Virginia Institute for Psychiatric and Behavior Genetics, Virginia Commonwealth University, VA, USA; QIMR Berghofer Medical Research Institute, Herston, Queensland 4006, Australia; Department of Psychiatry, University of California San Diego, La Jolla, CA, USA; Center for Behavior Genetics of Aging, University of California San Diego, La Jolla, CA, USA; Department of Psychological and Brain Sciences, Boston University, Boston MA, USA 02215; Department of Psychiatry and Behavioral Sciences, SUNY Upstate Medical University, Syracuse, NY, USA

**Author notes:** Corresponding author: Nathan A Gillespie. Drs. Kremen & Glatt are joint senior authors.

**Keywords:** weighted gene co-expression network analysis, module eigengenes, heritability

## Abstract

Multivariate network-based analytic methods such as weighted gene co-expression network analysis are being increasingly applied to human and animal gene-expression data to estimate module eigengenes (MEs). MEs represent multivariate summaries of correlated gene-expression patterns and network connectivity across genes within a module. Although this approach has the potential to elucidate the mechanisms by which molecular genomic variations contribute to individual differences in complex traits, the genetic etiology of MEs has never been empirically established. It is unclear if and to what degree individual differences in blood derived MEs reflect random variation versus familial aggregation arising from heritable or shared environmental influences. We used biometrical genetic analyses to estimate the contribution of genetic and environmental influences on MEs derived from blood lymphocytes collected on a sample of N=661 older male twins from the Vietnam Era Twin Study of Aging (VETSA) whose mean age at assessment was 67.7 years (SD=2.6 years, range=62-74 years). Of the 26 detected MEs, 14 (56%) had statistically significant additive genetic variation with an average heritability of 44% (SD=0.08, range=35-64%). Despite the relatively small sample size, this demonstration of significant family aggregation including estimates of heritability in 14 of the 26 MEs suggests that blood-based MEs are reliable and merit further exploration in terms of their associations with complex traits and diseases.

## Background

Gene expression (GE) is the first link in the chain connecting genetic polymorphisms to complex outcomes. GE is understood to play a critical role in many human diseases (Carroll; Emilsson et al., 2008; Kleinjan & Lettice, 2008; Kudaravalli et al., 2009; Schadt et al., 2005; Wray et al., 2007). Expression quantitative trait loci (eQTLs), which are single nucleotide polymorphisms (SNPs) regulating GE, provide a direct link between genome-wide association and GE studies (Franke & Jansen, 2009; Lehrmann & Freed, 2008). eQTL analysis can be useful for discerning transcriptome adaptations underlying complex behaviors (Bhattacharya & Mariani, 2009; Coppola & Geschwind, 2006; Gu et al., 2002). The identification of eQTLs located in transcription-factor-binding sites, splice sites, or other regulatory regions is likely to reveal mechanisms by which genetic variants contribute to individual differences in complex behaviors (Hindorff et al., 2009; Liu, 2011; Shastry, 2009). Yet, GE is also influenced by the environment, *via* epigenetic modifications. Thus, GE can be conceptualized as the primordial phenotype, and although likely influenced by genetic (G) and environmental (E) main effects as well as gene-by-environment interactions, the relative influence of genes and environment on GE is largely unknown.

A drawback of transcriptomic analysis is that functional genes with smaller effects on phenotypes may be undetectable at the single-gene level, particularly when relying on underpowered univariate statistics with high Type-I error rates (Langfelder & Horvath, 2008; Langfelder et al., 2013). Such analyses are also unlikely to capture biological interactions between genes, which seem likely to occur in complex traits (Horvath et al., 2006; Zhi et al., 2013). Indeed, notwithstanding questions regarding the direction of causation, dysregulation of disease-related gene networks is believed to be associated with many neurodevelopmental and psychiatric disorders (Oldham et al., 2008).

These limitations can be addressed with multivariate network-based analytic methods such as *weighted gene co-expression network analysis* (WGCNA) (Zhang & Horvath, 2005). WGCNA is a well-validated method for identifying genes associated with traits (DiLeo et al., 2011; Fuller et al., 2007; Langfelder & Horvath, 2008; MacLennan et al., 2009; Mason et al., 2009; Plaisier et al., 2009; Winden et al., 2011; Zhang & Horvath, 2005). It works by classifying a set of genes with correlated expression and connectivity patterns into ‘network modules’. Each of these networks is summarized via a single quantitative metric called a ‘*module eigengene’* (ME). Each ME is essentially the first principal component of a module, and as such, it summarizes some of the correlated expression across all the genes in the module. Yet, despite the increasing application of *MEs* in human and animal studies for exploring associations with complex behaviors and disorders (Chen et al., 2021; Liu et al., 2017; Wang et al., 2021; Zeng et al., 2020), the genetic and environmental etiology of MEs has never been empirically established.

For instance, it is unclear to what extent individual differences in MEs derived from blood lymphocyte transcriptomes are simply a reflection of random variation (including measurement error) versus familial aggregation attributable to either genetic or shared environmental influences. To address this gap, we used biometrical genetic analyses that rely on human twin data to explore the genetic and environmental variation in MEs derived from blood lymphocytes collected on a sample of older male twins.

## Materials & methods

### Subjects

Participants comprise middle-aged male twins from the Vietnam Era Twin Study of Aging (VETSA) (Kremen et al., 2013). The VETSA ascertained monozygotic and dizygotic twin-pairs who were concordant for US military service at some time between 1965-1975, although nearly 80% reported no combat experience. Based on data from the US National Center for Health Statistics, the sample is very similar to American men in their age cohort with respect to health and lifestyle characteristics (Schoeneborn & Heyman, 2009 ). The sample is 88.3% non-Hispanic white, 5.3% African-American, 3.4% Hispanic, and 3.0% “other” participants.

To date, VETSA twins have been assessed on three occasions, each 5-6 years apart. Briefly, Wave 1 took place between 2001 and 2007 (Kremen et al., 2006) (mean age=55.9, SD=2.4, range=51.1 to 60.7). Wave 2 occurred approximately 5.7 years later (mean age=61.7, SD=2.5, range=56.0 to 67.0). Wave 3 occurred a further 5.9 years later (mean age=67.6, SD=2.5, range=61.4 to 73.3). Data for this study came from a subset of Wave 3 subjects (N=695) who were approached to participate in a gene expression (GE) study. Here, the average age of subjects at the time of blood draw was 67.69 years (SD=2.6 years, range=62-74 years).

Subjects who participated in the GE study were marginally older than non-participants (67.4 vs 67.7, t = -2.03, df = 1157.20, p-value = 0.04).

### Ethics approval

Written informed consent was obtained from all participants. The University of California, San Diego (UCSD) and Boston University institutional review boards approved the proposal to collect these data (Projects 4013E, 150572, 150572, 150537, 140361, 071446, 031639, 151333). Data are publicly available through requests at the VETSA website (http://www.vetsatwins.org).

### Blood collection, RNA extraction & quality control

PAXgene tubes were used to collect 9ml of blood in total required for the RNA isolation. Total RNA was extracted from PAXgene tubes using whole blood gene RNA purification kits (QIAGEN). RNA from all samples was run on an Agilent Bioanalyzer to assess RNA quality and estimate RNA concentrations. Post-RNA quality assessment included total RNA conversion to cDNA, amplification, and purification using the Ambion Illumina TotalPrep RNA Amplification Kit (Ambion).

### RNA-seq analysis

RNA sequencing was performed on Illumina NovaSeq 6000 Sequencing System at the University of California, San Francisco and the UCSD IGM Genomics Center. Raw sequencing data in *Fastq* format (Babraham Bioinformatics, 2011) were transferred to SUNY Upstate Medical University and processed as follows.

Samples were sequenced on eight 96-well plates. An average 23 million paired-end 151 base pair reads per sample were obtained from each run. For quality control, three samples were duplicated on every plate. Sequencing quality was accessed by *fastQC* v0.11.8, with multiple quality statistics collected and evaluated at each processing step. One plate failed quality checks for 49 out of 96 samples and was discarded; all samples from this plate were re-sequenced as the eighth plate.

Low-quality bases and reads were filtered out of *Fastq* files by *trimmomatic* v0.39 (Bolger et al., 2014) with a quality score cutoff of 30. Trimmed reads were then mapped to the *Gencode* GRCh38.p12 release 30 human reference genome using *STAR aligner* v2.7.2b (Dobin et al., 2013). Mapping statistics were collected from alignment results using *PicardTools* v2.20.8 (Broad Institute, 2018). Reads were then summarized using *featureCounts* in *subread* v1.6.4 (Liao et al., 2014), and raw count data were combined and annotated using customized *R* scripts.

Transcripts, including protein-coding genes, miRNAs, and lncRNAs, were filtered to include only those with greater than or equal to one count per million in at least 50% of samples. Counts of zero were replaced with 0.5 and data were normalized to effective library size via the *TMM* method in *edgeR* v3.28.1(Robinson et al., 2010). Batch effects were reduced using *ComBat* (Wang et al., 2017).

### Weighted gene co-expression network analysis (WGCNA)

We performed weighted gene co-expression network analysis (WGCNA) (Zhang & Horvath, 2005) on the GE data to identify the GE networks using the *WGCNA* v1.36 package in *R* (Langfelder & Horvath, 2008; Langfelder et al., 2013; Zhang & Horvath, 2005). This approach identifies higher-order interactions between genes by assembling the GE data into ‘network modules’. Each network module in WGCNA was then represented using a ‘module eigengene’ (ME), which is a quantitative summary of the correlated expression and connectedness across all the genes within each module.

The *goodsamplesgenes*() function identifies genes and samples with low variance or too much missingness, and excludes these from network construction. This function was run to ensure suitability of the features and subjects, detecting no samples or genes to be removed. Both functions relied on default parameters. The *blockwiseModules*() function was then used to create a signed coexpression network. Soft threshold power was set at 12 following the developer’s recommendation (Langfelder & Horvath) *deepSplit* was set to 2, with a minimum module size of 30. Using these parameters, expression data for 4571 transcripts were organized into 26 modules with sizes ranging from 114 to 1,200 transcripts. The 26 modules include a ‘grey’ or unclassified bin containing genes not reliably connected (ME0).

### Residualization & descriptive statistics

MEs were generated for 692 VETSA subjects. However, to improve the distribution, all ME means and variances were residualized for the effects of testing site, time between blood draw and storage, and storage times at -20 C and -80 C using the *umx_residualize* function (version 4.9.0) in R (version 4.1.1). A total of 31 cases were lost during residualization due to missing covariates. Another two subjects were removed due to missing zygosity. Descriptive statistics for each ME following residualization are shown in Table 1. Among the 661 individuals with complete MEs there were 160 monozygotic twin pairs, 67 monozygotic singletons (incomplete pairs), 100 dizygotic twin pairs, and 74 dizygotic singletons.

**Table 1.** Descriptive statistics along with monozygotic (MZ) and dizygotic (DZ) twin pair polychoric correlations and their 95% confidence intervals for each module eigengene (ME).

|  | N | Mean | SD | Minimum | Maximum | MZ | (95% CI) | DZ | (95% CI) |
| --- | --- | --- | --- | --- | --- | --- | --- | --- | --- |
| ME1 | 661 | 0.0 | 0.0279 | -0.1118 | 0.1279 | 0.49 | (0.35, 0.60) | 0.09 | (-0.13, 0.30) |
| ME2 | 661 | 0.0 | 0.0281 | -0.0967 | 0.1246 | 0.43 | (0.28, 0.55) | 0.05 | (-0.17, 0.26) |
| ME3 | 661 | 0.0 | 0.0346 | -0.1549 | 0.1206 | 0.38 | (0.22, 0.51) | 0.14 | (-0.06, 0.33) |
| ME4 | 661 | 0.0 | 0.0270 | -0.1050 | 0.0775 | 0.46 | (0.33, 0.58) | 0.36 | (0.15, 0.53) |
| ME5 | 661 | 0.0 | 0.0337 | -0.1263 | 0.1239 | 0.38 | (0.24, 0.51) | 0.13 | (-0.07, 0.32) |
| ME6 | 661 | 0.0 | 0.0338 | -0.1600 | 0.1050 | 0.38 | (0.23, 0.51) | 0.29 | (0.09, 0.47) |
| ME7 | 661 | 0.0 | 0.0338 | -0.1080 | 0.1448 | 0.36 | (0.22, 0.49) | 0.13 | (-0.09, 0.33) |
| ME8 | 661 | 0.0 | 0.0374 | -0.0940 | 0.1497 | 0.42 | (0.28, 0.54) | 0.30 | (0.10, 0.47) |
| ME9 | 661 | 0.0 | 0.0349 | -0.1151 | 0.1405 | 0.42 | (0.28, 0.55) | 0.15 | (-0.06, 0.35) |
| ME10 | 661 | 0.0 | 0.0326 | -0.1080 | 0.1130 | 0.45 | (0.31, 0.56) | 0.36 | (0.18, 0.52) |
| ME11 | 661 | 0.0 | 0.0368 | -0.1403 | 0.1001 | 0.40 | (0.25, 0.52) | 0.35 | (0.17, 0.51) |
| ME12 | 661 | 0.0 | 0.0291 | -0.1460 | 0.1178 | 0.51 | (0.38, 0.61) | 0.27 | (0.07, 0.44) |
| ME13 | 661 | 0.0 | 0.0300 | -0.1281 | 0.0767 | 0.58 | (0.46, 0.67) | 0.47 | (0.30, 0.60) |
| ME14 | 661 | 0.0 | 0.0357 | -0.1452 | 0.1035 | 0.40 | (0.25, 0.53) | 0.30 | (0.11, 0.47) |
| ME15 | 661 | 0.0 | 0.0364 | -0.1144 | 0.1291 | 0.42 | (0.27, 0.54) | 0.14 | (-0.07, 0.33) |
| ME16 | 661 | 0.0 | 0.0365 | -0.1186 | 0.1139 | 0.50 | (0.38, 0.60) | 0.19 | (-0.01, 0.37) |
| ME17 | 661 | 0.0 | 0.0371 | -0.1369 | 0.1247 | 0.27 | (0.11, 0.41) | 0.26 | (0.04, 0.45) |
| ME18 | 661 | 0.0 | 0.0370 | -0.1045 | 0.1114 | 0.59 | (0.48, 0.68) | 0.26 | (0.06, 0.43) |
| ME19 | 661 | 0.0 | 0.0380 | -0.1307 | 0.1418 | 0.50 | (0.37, 0.61) | 0.27 | (0.07, 0.45) |
| ME20 | 661 | 0.0 | 0.0342 | -0.1052 | 0.1267 | 0.44 | (0.31, 0.56) | 0.15 | (-0.06, 0.34) |
| ME21 | 661 | 0.0 | 0.0359 | -0.1214 | 0.1208 | 0.48 | (0.35, 0.59) | 0.22 | (0.02, 0.40) |
| ME22 | 661 | 0.0 | 0.0353 | -0.1135 | 0.1380 | 0.52 | (0.40, 0.62) | 0.03 | (-0.17, 0.23) |
| ME23 | 661 | 0.0 | 0.0365 | -0.0642 | 0.1998 | 0.34 | (0.20, 0.46) | 0.12 | (-0.05, 0.29) |
| ME24 | 661 | 0.0 | 0.0365 | -0.0933 | 0.1368 | 0.39 | (0.26, 0.51) | 0.10 | (-0.16, 0.34) |
| ME25 | 661 | 0.0 | 0.0367 | -0.0733 | 0.1187 | 0.65 | (0.55, 0.73) | 0.25 | (0.06, 0.41) |
| ME0 | 661 | 0.0 | 0.0351 | -0.1283 | 0.1258 | 0.38 | (0.24, 0.51) | 0.33 | (0.13, 0.50) |

Footnotes: Residualization was performed using the `umx_residualize` function (version 4.9.0) in R (version 4.1.1). All MEs were residualized for i) testing site, ii) time in hours from blood draw to storage in -20 C, iii) time in days spent in -20 C storage, iv) time in days spent in $\pm 80$ C storage, v) time in days between blood draw & RNA extraction, & vi) time in days between blood draw and sequencing. A total of 32 cases were lost due to missing covariates.

### Statistical analyses

The *OpenMx*_2.9.9.1_ software package (Boker et al., 2011) in *R*_3.4.1_ (R Development Core Team, 2018) was used to fit univariate biometric genetic twin models (Neale & Cardon, 1992) to estimate the relative contribution of genetic and environmental influences in each ME. Given the number of incomplete twin pairs, methods such as Weighted Least Squares will result in significant listwise deletion thereby altering the accuracy of each ME’s mean and variance.

Fortunately, the raw data Full Information Maximum Likelihood (FIML) option in *OpenMx*_2.9.9.1_ (Boker et al., 2011) is robust to violations of non-normality while enabling analysis of missing or incomplete data as well as the direct estimation of covariate effects. We included age at blood draw as a covariate. This enables accurate estimates of means thereby improving estimation of the variances and covariance structure used to test our competing hypotheses regarding the genetic and environmental etiology of each ME.

### Univariate analyses

In univariate analyses, the total variation in each of the 26 MEs was decomposed into additive (A) genetic (heritability), shared or common environmental (C), and non-shared or unique (E) environmental variance components (see Figure 1). This approach is referred to as an ‘ACE’ variance component model. The decomposition is achieved by exploiting the expected genetic and environmental correlations between monozygotic (MZ) and dizygotic (DZ) twin pairs. MZ twin pairs are genetically identical, whereas DZ twin pairs share, on average, half of their genes. Therefore, the MZ and DZ twin pair correlations for the additive genetic effects are fixed to r_A_=1.0 and r_A_=0.5 respectively. The modelling assumes that shared environmental effects (C) are equal in MZ and DZ twin pairs (r_C_=1.0), while non-shared environmental effects (E) are by definition uncorrelated. Shared environmental effects are defined as those that make siblings similar. Non-shared environmental factors are defined as those that make siblings different; this term also includes measurement error. Note that these variance components are derived from the association between phenotypic and genotypic similarities of the twin pairs.

**Figure 1.**
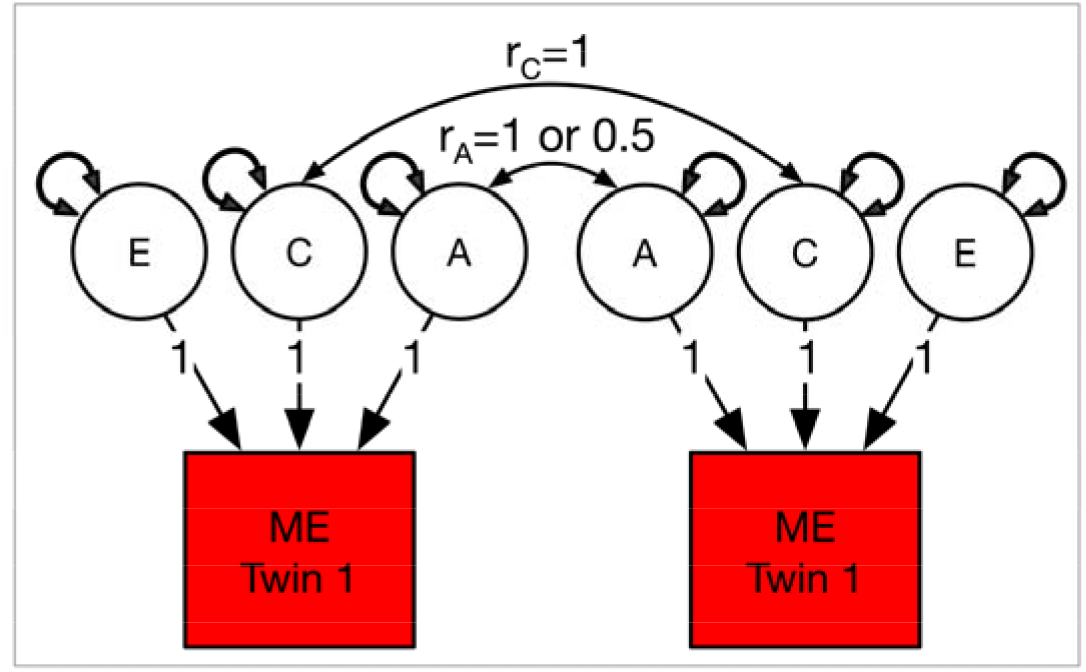
Univariate variance decomposition to estimate the relative contribution of genetic & environmental influences in each module eigengene (ME). Footnotes: A = additive genetic, C = common or shared environmental, & E = unshared environmental influences. Single-headed arrows represent the causal contributions of the A, C & E effect for a given ME. Double-headed arrows represent A, C, & E latent factor variances & twin pair covariances. r_C_ = correlation of 1 for MZ and DZ twin pairs. r_A_ = 1 or 0.5 for MZ & DZ twin-pairs respectively.

### Model fit

For each univariate analysis, we fitted an A+C+E model before determining the most likely sources of variance by fitting three additional sub-models in which the i) A, ii) C, and iii) A and C influences were fixed to zero. In other words, the significance of each nested sub-model (i.e. ‘AE’, ‘CE’ and ‘E’) was determined using the change in minus twice the Log-Likelihood (Δ-2LL) when compared to the full ACE model. Under certain regularity conditions, this Δ-2LL is asymptotically distributed as chi-squared with the degrees of freedom equal to the difference in the number of free parameters in the two models (Steiger et al., 1985). Our determination of the best-fitting model was also based on the optimal balance of complexity and explanatory power by selecting the model with the lowest Akaike’s Information Criterion (AIC) value (Akaike, 1987).

## Results

### Twin pair correlations

Table 1 shows the twin pair correlations by zygosity for all 26 MEs. If familial aggregation is entirely attributable to shared environmental influences, then the MZ and DZ twin pair correlations will be statistically equal. If, however, familial aggregation is attributable to shared additive (or non-additive) genetic factors, then DZ correlations will be less than ½ the MZ twin pair correlations. Finally, if the hypothesis is that blood-based MEs are unreliable or that variability arises entirely from either random influences that are unshared between siblings or from measurement error, then both MZ and DZ twin pair correlations will not statistically differ from zero. For ME4, ME6, ME8, MEs10-11, MEs13-14, ME17 and ME0, the DZ twin pair correlations were greater than ½ of their MZ twin pair counterparts, which is consistent with familial aggregation attributable to shared or common environmental (C) influences. In contrasts, DZ twin pair correlations for MEs ME3, ME7, ME12, MEs18-19, ME21 and ME23 were approximately ½ (±0.05) of their MZ twin pair counterparts, which is consistent with familial aggregation driven by additive genetic influences. For the remaining MEs (ME1-2, ME9, MEs15-ME16, ME20, ME22, and MEs24-25), the DZ twin pair correlations were well below their MZ twin pair counterparts, which is consistent with non-additivity and dominance (D) influencing familial aggregation. In the Classical Twin Design, ‘C’ and ‘D’ influences are negatively confounded, and therefore, cannot be modelled simultaneously (Martin et al., 1978). Since the sample sizes required to detect ‘D’ as a source of variation are very large, even for variables measured on a continuous liability scale, we chose to model ‘C’ influences in all subsequent univariate and multivariate models below.

### Twin modeling

Supplementary Table S1 shows the full univariate model fitting results, including comparisons between the ‘ACE’ and the ‘AE’, ‘CE’ and ‘E’ sub-models. Table S1 also includes 95% confidence intervals for all A, C and E variance components along with 95% confidence intervals. Among the 26 MEs examined, there were 15 MEs with evidence of significant familial aggregation attributable to either additive (A) genetic or common (C) environmental influences (see Table 2).

**Table 2.** Standardized estimates (including 95% confidence intervals [95%CI]) of additive genetic (A), common environment (C), & non-shared environment (E) influences under the best fitting univariate models for each module eigengene (ME).

| **MEs with significant additive genetic variation** |  |  |  |  |  |  |
| --- | --- | --- | --- | --- | --- | --- |
|  | **A** | **95%CI** | **C** | **95%CI** | **E** | **95%CI** |
| ME1 | 0.44 | (0.31, 0.54) | - | - | 0.56 | (0.46, 0.69) |
| ME2 | 0.38 | (0.24, 0.49) | - | - | 0.62 | (0.51, 0.76) |
| ME5 | 0.38 | (0.24, 0.50) | - | - | 0.62 | (0.50, 0.76) |
| ME7 | 0.36 | (0.22, 0.48) | - | - | 0.64 | (0.52, 0.78) |
| ME9 | 0.40 | (0.27, 0.52) | - | - | 0.60 | (0.48, 0.73) |
| ME16 | 0.51 | (0.39, 0.61) | - | - | 0.49 | (0.39, 0.61) |
| ME18 | 0.58 | (0.48, 0.67) | - | - | 0.42 | (0.33, 0.52) |
| ME19 | 0.52 | (0.40, 0.62) | - | - | 0.48 | (0.38, 0.60) |
| ME20 | 0.41 | (0.29, 0.52) | - | - | 0.59 | (0.53, 0.71) |
| ME21 | 0.47 | (0.35, 0.57) | - | - | 0.53 | (0.43, 0.65) |
| ME22 | 0.50 | (0.37, 0.61) | - | - | 0.50 | (0.39, 0.63) |
| ME23 | 0.35 | (0.21, 0.47) | - | - | 0.65 | (0.53, 0.79) |
| ME24 | 0.38 | (0.25, 0.49) | - | - | 0.62 | (0.51, 0.75) |
| ME25 | 0.64 | (0.54, 0.71) | - | - | 0.36 | (0.29, 0.46) |

| **MEs with significant common environment variation** |  |  |  |  |  |  |
| --- | --- | --- | --- | --- | --- | --- |
|  | **A** | **95%CI** | **C** | **95%CI** | **E** | **95%CI** |
| ME10 | - | - | 0.42 | (0.31, 0.51) | 0.58 | (0.49, 0.69) |
| ME13 | - | - | 0.54 | (0.45, 0.62) | 0.46 | (0.38, 0.55) |

| **MEs with no significant familial aggregation** |  |  |  |  |  |  |
| --- | --- | --- | --- | --- | --- | --- |
|  | **A** | **95%CI** | **C** | **95%CI** | **E** | **95%CI** |
| ME3 | 0.39 | (-0.09, 0.91) | -0.04 | (-0.50, 0.37) | 0.65 | (0.52, 0.79) |
| ME4 | 0.09 | (-0.31, 1.00) | 0.35 | (-0.10, 0.69) | 0.56 | (0.45, 0.68) |
| ME6 | 0.18 | (-0.26, 0.66) | 0.20 | (-0.23, 0.56) | 0.62 | (0.50, 0.77) |
| ME8 | 0.27 | (-0.14, 0.72) | 0.15 | (-0.25, 0.50) | 0.57 | (0.46, 0.71) |
| ME11 | 0.14 | (-0.26, 0.57) | 0.27 | (-0.11, 0.59) | 0.59 | (0.47, 0.73) |
| ME12 | 0.33 | (-0.08, 0.81) | 0.15 | (-0.31, 0.50) | 0.52 | (0.42, 0.64) |
| ME14 | 0.15 | (-0.29, 0.62) | 0.24 | (-0.18, 0.59) | 0.61 | (0.49, 0.76) |
| ME15 | 0.46 | (-0.03, 0.99) | -0.07 | (-0.56, 0.36) | 0.61 | (0.50, 0.74) |
| ME17 | 0.15 | (-0.33, 0.63) | 0.15 | (-0.26, 0.52) | 0.70 | (0.56, 0.87) |
| ME0 | -0.02 | (-0.44, 0.46) | 0.38 | (-0.06, 0.73) | 0.64 | (0.52, 0.77) |

### MEs with significant additive genetic familial aggregation

There were 14 MEs (ME1, ME2, ME5, ME7, ME9, ME16, & MEs18-25) in which the 95% confidence intervals surrounding the additive genetic ‘A’ point estimates in the saturated ACE model did not include zero (Supplementary Table S1). For these MEs, dropping the ‘A’ influences resulted in a significant change in chi-square, whereas removing the common ‘C’ environmental influences (i.e. the AE model) improved the model fit in terms of a non-significant change in chi-square and producing the lowest AIC value. As illustrated in Figure 2, additive genetic influences explained 35% to 64% of the total variance across these 14 MEs (mean = 45%, median = 43%, standard deviation = 9%).

**Figure 2.**
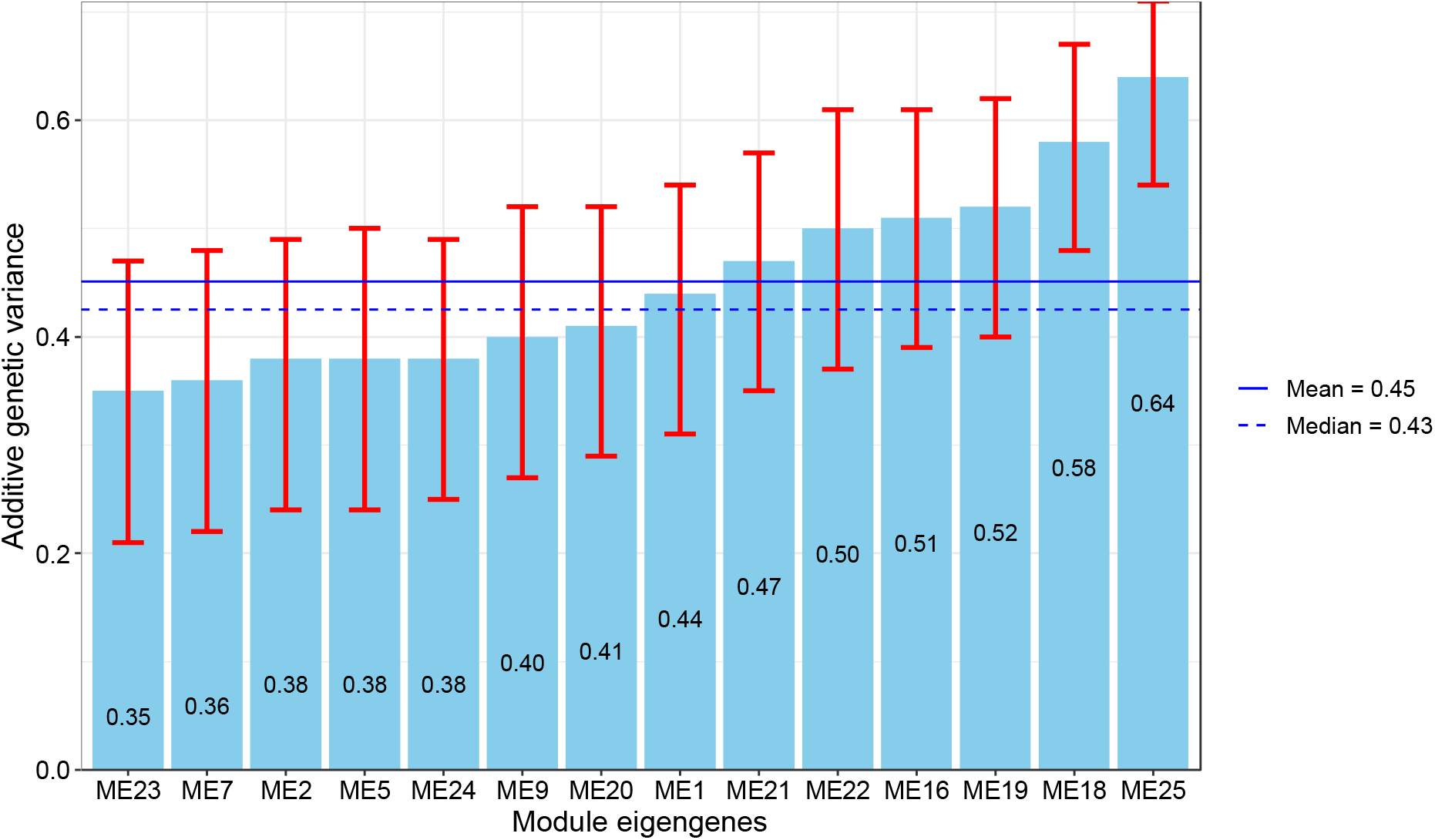
Standardized estimates (including 95% confidence intervals) of additive genetic variance in ascending order for the 14 MEs with significant heritability.

### MEs with significant common environment familial aggregation

There were two MEs (ME10 & ME13) in which the 95% confidence intervals surrounding the ‘C’ point estimates in the saturated ACE univariate model did not include zero. In both cases, dropping the ‘C’ influences resulted in a significant change in chi-square, whereas removing the ‘A’ influences (i.e. the CE model) improved the model fit in terms of a non-significant change in chi-square. Common environmental influences explained 42% to 54% of the total variances in ME10 and ME13 respectively.

In small twin studies, estimates of ‘C’ should always be interpreted in the context of statistical power. We, therefore, performed post-hoc power calculations using the *umx* package *power*.*ACE*.*test* (Bates et al., 2019) to determine the minimum amount of ‘C’ that could be detected at 80% power (see Supplement). Given the current sample size, we estimated that there was 80% power to detect estimates of ‘C’ as low as 44%. In terms of the power to detect the ‘C’ estimate of 42% observed in ME10, the power was 73%.

### MEs without significant familial aggregation

For the remaining 11 MEs (ME3, ME4, ME6, ME8, ME11-12, ME14-15, ME17 & ME0) there was insufficient power to choose between the competing ‘AE’ and ‘CE’ univariate models.

Consequently, we retained the saturated ‘ACE’ model as the best fitting in each instance. Note that in each of these MEs, the 95% confidence intervals surrounding both the ‘A’ *and* ‘C’ point estimates spanned zero.

## Discussion

To our knowledge, this is the first study to explore the genetic and environmental contributions to weighted gene co-expression network analysis module eigengenes (MEs). Among the 26 derived MEs, 50% had statistically significant additive genetic variation with an average heritability of 46% (SD=0.09, range=35-64%). Additionally, there were two MEs with evidence of significant shared environmental influences accounting for 42% to 54% of the total variance. For the remaining 11 MEs, estimates of additive genetic *and* shared environmental influences were non-significantly different from zero. Despite the small sample size for a classical twin study, the demonstration of significant family aggregation including estimates of significant heritability suggests that blood-based gene expression networks are reliable and merit further exploration in terms of their associations with complex traits and diseases.

Our average estimate of ME heritability is higher than the heritability estimates of individual differentially expressed genes reported previously. For example, Monks et al.’s analyses of human lymphoblastoid cell lines (LCLs) taken from 167 individuals found that among N=2,340 differentially expressed genes, 31% (N=762) were significantly heritable (*h*^2^), with a median *h*^2^ of 34% (Monks et al., 2004). McRae et al. estimated a mean heritability of 31% for normalized gene expression levels based on N=15,887 transcripts in LCLs taken from a small sample of monozygotic twin pairs (MZ) (McRae et al., 2007). Powell et al. (Powell et al., 2012) analysed 9,555 genes in LCLs and whole blood (WB) taken from 50 and 47 MZ twin pairs, and estimated average heritabilities for differential gene expression of 38% for LCLs and 32% for WB. We note that the methods used by these three reports captured broad sense heritability and assumed an absence of common environmental influences; our results have shown these ‘C’ influences to be present in at least two module eigengenes. Wright et al. (Wright et al., 2014) used the classical twin design, which unconfounds additive genetic and common environmental influences, to analyze gene expression levels based on peripheral venous blood taken from 2,752 twins. Among differentially expressed genes, they reported an average heritability of 14% (±15%). While these previous reports demonstrate a clear pattern of heritability for QTLs, our findings suggest that the average heritability of MEs is potentially higher than differentially expressed genes. Complex gene networks and interactions are the hallmark of complex traits (Horvath et al., 2006; Zhi et al., 2013), and since MEs are multivariate summaries of correlated gene expression patterns and network connectivity across all genes in a module, the signal detection power when based on the heritable MEs is likely to be greater compared to analyses based on univariate GE loci (Horvath, 2011).

Of interest are the two MEs with evidence of significant shared environmental (C) influences. As with additive genetics, ‘C’ influences also drive twin pair similarity. The advantage of the twin methodology applied here is that the use of age-matched siblings reared together provides a natural control for environmental influences that were either shared during infancy and youth, continue to be shared, or are persistent and continue to exert an impact. Naturally, as siblings age and spend less time together, the number of directly shared environmental influences, relative to the time in their lives when reared together, is expected to diminish. Despite the expected waning of environmental influences shared between siblings, familial aggregation in ME10 & ME13 was unequivocally explained by shared environmental influences ranging from 42% to 54% of the total variance. Given that batch effects can also drive twin pair similarity when correlated within twin pairs, all 26 MEs were corrected for the effects of testing location, the time from blood draw to placement in -20 C storage, total time spent in -20 C storage and Ill80 C storage, the time between blood draw and RNA extraction, and finally, the time between blood draw and sequencing. Although caution should normally be applied when interpreting ‘C’ in small twin studies, the power to detect the ‘C’ influence of 41% for ME10 in this sample was 73%. If, after future efforts at validation and replication based on larger samples, estimates of C persist, then further analyses should elucidate which aspects of environments shared between siblings, broadly indexed by ‘C’ here, are responsible for these observed individual differences e.g., allostatic stressors, diet, geographical location, etc.

Our results must be interpreted in the context of at least two limitations. First, between-group differences are likely to exist and may be complex (Royse et al., 2021). Therefore, our results may not generalize to women or other age or ancestral groups. It is unclear to what extent there are sex differences in the genetic and environmental etiology of MEs in similarly aged women. It is plausible that the genetic and environmental etiologies of these MEs may differ between ancestral groups. Only by ascertaining larger and ancestrally varied samples can we begin to test hypotheses regarding sex and ancestral group differences.

Second, while estimating GE patterns from whole blood has become a common alternative to *postmortem* brain studies (GTEx Consortium, 2013, 2015), our analyses did not account for cell heterogeneity, which can bias results. For instance, cell heterogeneity can result in GE differences due to different proportions of cells between samples, or mask important disease-relevant differences. It is therefore important to control for such differences in order to extract accurate and meaningful GE information. One solution would be to account for cell-specific contributions is assessing GE patterns in certain cell populations derived from fluorescence-activated cell sorting (FACS) approaches. An alternative, emerging solution would be to rely on ‘*computational deconvolution’* methods to infer cell type-specific information from data generated from heterogeneous samples (Novershtern et al., 2011; Shay & Kang, 2013; Shen-Orr & Gaujoux, 2013).

## Conclusion

This is first study to explore the genetic and environmental contributions to weighted gene co-expression network analysis module eigengenes (MEs). Among 14 MEs with statistically significant additive genetic variation, the average heritability was 46%. Because network modules may not be well preserved between blood and brain (Cai et al., 2010), blood GE may not recapitulate neurobiological mechanisms of complex behaviors and disorders. Therefore, while blood GE and MEs can be used as proxy biomarkers for brain-based phenotypes and complex traits, blood tissue should only be considered a useful proxy for gene expression in the brain provided that the relevant gene(s) are expressed in both tissues (Joehanes et al., 2012; Sullivan et al., 2006). To address this problem, we have recently developed a novel transcriptome-imputation method called Brain Gene Expression and Network Imputation Engine (BrainGENIE) (Hess et al., 2023) that is capable of imputing brain-region-specific gene expression levels from peripheral blood gene expression. In addition to applying BrainGENIE to the current data, it remains an empirical question as to how well blood-based MEs assessed in early old age can be leveraged as useful biomarkers for complex traits and disorders that emerge later in life.

## Acknowledgments

We kindly thank the twins from the Vietnam Era Twin Study of Aging (VETSA) for their participation.

