## Supplement for "Genetic and environmental contributions to eigengene expression"

*Post-hoc power calculations*

Post-hoc power estimation was conducted using the umx (Bates et al., 2019) option power.ACE.test() to determine the minimum amount of shared environmental influences that can be detected at 80% given the current sample. The power.ACE.test() parameters included N=663/2 twin-pairs, an MZ-DZ sampling ratio of 1.4, with the method option set to ‘empirical’. There was 80% to detect a minimum of ~43% shared environmental influences.
